# Identifying Taxonomic Units in Metagenomic DNA Streams

**DOI:** 10.1101/2020.08.21.261313

**Authors:** Vicky Zheng, Ahmet Erdem Sariyuce, Jaroslaw Zola

## Abstract

With the emergence of portable DNA sequencers, such as Oxford Nanopore Technology MinION, metagenomic DNA sequencing can be performed in real-time and directly in the field. However, because metagenomic DNA analysis is computationally and memory intensive, and the current methods are designed for batch processing, the current metagenomic tools are not well suited for mobile devices.

In this paper, we propose a new memory-efficient method to identify Operational Taxonomic Units (OTUs) in metagenomic DNA streams. Our method is based on finding connected components in overlap graphs constructed over a real-time stream of long DNA reads as produced by MinION platform. We propose an efficient algorithm to maintain connected components when an overlap graph is streamed, and show how redundant information can be removed from the stream by transitive closures. Through experiments on simulated and real-world metagenomic data, we demonstrate that the resulting solution is able to recover OTUs with high precision while remaining suitable for mobile computing devices.

## 1 Introduction

Recently introduced nanopore-based DNA sequencers, specifically Oxford Nanopore Technology (ONT) MinION [30], are revolutionizing how DNA-based studies are performed. Their key advantages are a small form factor and low energy consumption that make them fully portable and allow for easy deployment in the field, outside of a typical laboratory [16, 31]. Moreover, these devices can sequence DNA molecules directly (i.e., without extra steps like DNA amplification), and can stream the resulting *reads,* which are strings of A,C,G,Ts, in near real-time. This makes them extremely attractive for *metagenomic* studies that involve processing DNA recovered directly from environmental samples. In recent years, MinION sequencers have been increasingly used for *in situ* studies, including, for example, tracking of COVID19, Ebola and Zika outbreaks [8, 22, 27], deployments in the Arctic and Antarctic [7], and even on the International Space Station [5] (we invite reader to [13] for a broader discussion on mobile DNA sequencing).

One of the most common tasks in metagenomics is identification of Operational Taxonomic Units (OTUs) represented by clusters of highly similar DNA reads. OTUs often serve as a proxy representing microbial composition of the sequenced sample in cases where reads classification (e.g., by searching a DNA database of known organisms) is difficult or impossible. However, in the current mobile DNA sequencing workflows, identification of OTUs, along with any other DNA analytics, remains challenging. In Figure 1, we outline the usual mobile DNA sequencing workflow using MinION. The device streams, in real-time, electric signals characterizing detected DNA fragments. These signals are basecalled to yield DNA reads, which are next processed using full-fledged bioinformatics tools. In a mobile setup, the sequencer is typically coupled with a portable host device with limited compute power, memory, and energy supply (e.g., tablet or a dedicated system-on-a-chip like for example MinIT [18]). Since the basecalling process is already memory and compute intensive, the bioinformatics analysis step has to be either offloaded to a cloud service (which is not always possible or desired), or postponed until sufficient compute resources become available. In both cases, the resulting delay between DNA read acquisition and the analytics is highly undesired from the end-user’s perspective, as it decreases the overall responsiveness.

**Figure 1:**
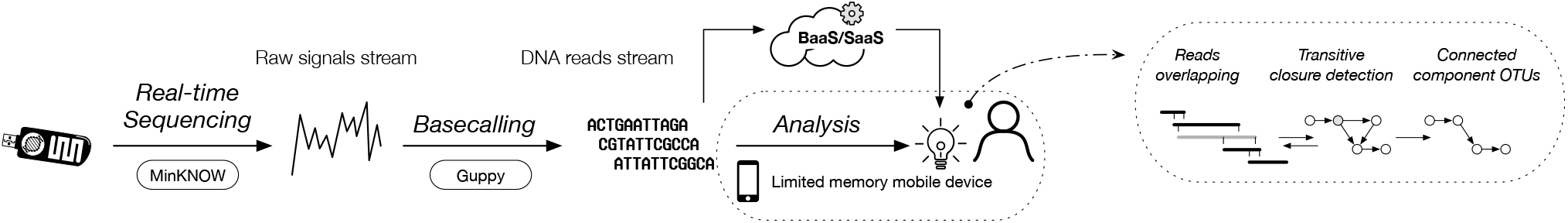
Schematic representation of mobileDNAsequencing pipeline with MinION. Sequenced reads can be processed locally, or can be offload to a Back-end as a Service (or Software as a Service) in a cloud. In this work, we aim at OTUs identification method suitable to run directly on mobile devices.

In this paper, we focus on the problem of identifying OTUs in mobile DNA read streams generated by MinION portable sequencers. Our goal is to provide memory and compute efficient solution that could be deployed as a co-processing routine in portable DNA sequencing workflows operating on light-weight computational devices. Our approach is based on finding connected components in the similarity (or overlap) graphs constructed and streamed directly over the DNA reads streams. Connected components have been demonstrated as a robust representation of OTUs [9, 10, 25], and are very attractive abstraction, as they are starting point to multiple other tasks, including DNA assembly [26], reference-free taxonomic classification (or clustering) [12, 34], or species abundance estimation [6]. In the paper, we first propose efficient algorithms to identify transitive closures and maintain connected components in streaming graphs. We then show how our method can be coupled with DNA overlapper routine to handle real-world DNA processing workloads without sacrificing performance. Through experimental results we demonstrate that the proposed method is memory efficient, and can identify OTUs in ONT MinION sequencing data.

The remainder of this paper is organized in the usual way: in Section 2, we formalize the problem and introduce our notation. In Section 3, we give detailed description of our approach. We follow with experimental validation and discussion in Section 4. We close the paper with a brief review of related work in Section 5, and conclusions in Section 7.

## 2 Problem Formulation

We consider mobile DNA sequencing pipeline as presented in Figure 1. Portable DNA sequencer (specifically ONT MinION) is attached and controlled by a battery powered mobile host (e.g., laptop) that is also responsible for DNA data processing. The sequencer delivers, in real-time, raw signals representing detected DNA molecules. These raw signals are immediately basecalled on the host machine yielding the actual DNA reads. To illustrate the rate at which the process happens: in our experiments, we usually observe that the sequencer delivers around 31 raw signals per second. The basecalling rate varies depending on the host machine capabilities. For example, using NVIDIA Nano Supercomputer-on-a-Chip with 128 GPGPU cores and 4GB of main memory, the basecalling can be sustained at the rate of approximately 130 reads per second. The resulting DNA reads stream is passed for the downstream analysis. In this work, we are specifically interested in performing OTUs identification directly on the host that receives streamed DNA reads.

To formalize our problem, let *R* = [*r*_1_,…, *r*_*n*_] represent an input stream of DNA reads generated in real-time by a DNA sequencer. We have that for each *i* < *j* read r; precedes read *r*_*j*_ in the stream, which we will denote by *r*_*i*_ ≺ *r*_*j*_. The size of the stream, n, is not known *a priori.* For example, a user may decide to terminate sequencing experiment at any point of time (e.g., after *sufficient* data has been collected), or may run experiment for a specified time interval (e.g., 2h, which could be a small metagenomic experiment).

Given a set of DNA reads, we can construct an overlap graph *G* = (*V*, *E*) in which vertices *V* represent the reads, and two vertices, u and v, are connected by the directed edge, *u* → *v*, denoted by *e* = (*u*, *v*), if there is a *significant* overlap between suffix of the read represented by u and prefix of the read represented by v. Here significant overlap means that the length of the suffix-prefix match is beyond some predefined threshold, and indicates that the two reads corresponding to u and v have been derived from neighboring portions of the unknown underlying genome. For now, we assume that an overlapper software is available and capable to construct overlap graph over the stream R (see below).

Let R^*i*^ be the set of first *i* reads from the stream *R*, and *G*^*i*^ = *(V*^*i*^*, E*^*i*^ ) be the overlap graph constructed over *R*^*i*^. Moreover, let 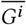 denote an undirected graph constructed by treating all edges in E^*i*^ as undirected. Our goal is to dynamically identify and maintain, in a computationally and memory efficient way, a set 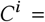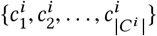 of all connected components found in graph 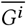, where 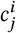 is a set of vertices in component *j* of graph 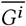. The final set of connected components, *C*^*n*^, will represent the Operational Taxonomic Units over the set of DNA reads in *R*, i.e., we will expect that all reads within a given connected component will be coming from the same taxonomic unit (e.g., an organism). At this point we should note that,in our formulation, we use connected components as proxy for OTUs - as mentioned earlier, this approach has been demonstrated as a robust strategy in the past [9, 10, 25] and hence we adopt it here.

We note that when processing stream R we are concerned only with graph *G*^*i*^ and set *C*^*i*^ (since these structures carry our information of interest), and do not have to explicitly store or maintain graph 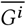. Moreover, currently we are not concerned with the details of how the overlap graph is computed (e.g., what is the similarity threshold for suffix-prefix comparison, or how DNA reverse-complements are handled, we discuss this issue in Section 3.4). In other words, we are assuming that in our mobile DNA sequencing pipeline, there is a read overlapper tool that operates under the hardware constraints mentioned earlier. The overlapper handles incoming stream of reads generated by a sequencer, and creates, with some precision and sensitivity, a new stream of edges induced by the incoming read. Specifically, given incoming read *r*_*i*_, let *v*_*i*_ be its corresponding vertex in graph *G*^*i*^. The overlapper provides sets *N*^+^(*v*_*i*_) ⊆ *V*^*i*−1^ and *N^−^(v_*i*_)* ⊆ V^*i*−1^, such that for each *u* ∊ *N*^+^(*v*_*i*_) there is an edge (*v*_*i*_, *u*) ∊ *E*^*i*^ and for each *u* ∊ *N*^−^(*v*_*i*_) there exists an edge (*u*, *v*_*i*_) ∊ *E*^*i*^ (note that in both cases u corresponds to a read that precedes *r*_*i*_ in *R*).

## 3 Proposed Approach

Given the above problem formulation, to create and maintain our desired set *C*^*i*^, we could leverage one of the several existing algorithms for finding connected components in graph streams (we review them in Section 5). However, this approach has drawbacks since the current algorithms are tailored for general graph streams and are oblivious to the domain specific properties of the data. Therefore in our approach we take a slightly different route and exploit the (well known) fact that, in a typical DNA sequencing experiment, many DNA reads are sharing an overlap, and thus provide redundant information.

To better illustrate this property, consider a set of reads in Figure 2a that come from the same region of a genome. Because read *r*_*j*_ overlaps with both *r*_*i*_ and *r*_*k*_, and at the same time *r*_*i*_ and *r*_*k*_ are overlapping as well, we can eliminate read *r*_*j*_ without disconnecting the underlying overlap graph and hence without losing any critical information. This simplifies the graph on which we have to perform connected components identification, and reduces the number of reads (and hence graph nodes) we have to maintain in the memory. The redundant reads may be marked as such, and can be offloaded to a persistent storage and processed later, e.g., when more computational resources are available.

**Figure 2:**
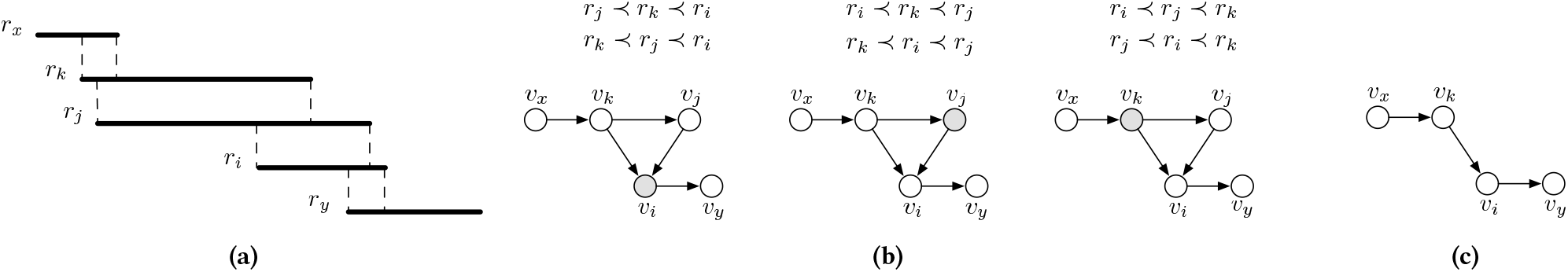
(a) Example DNA reads with their overlaps marked, (b) Three different cases that may occur in the overlap graph constructed over reads in (a) if *r*_*x*_ ≺ *r*_*y*_ ≺ *r*_*i*_, *r*_*j*_, *r*_*k*_, and overlapper does not report overlap between *r*_*x*_ → *r*_*j*_ and *r*_*j*_ → *r*_*y*_ (c) Irreducible graph created by removing *r*_*j*_ as a transitive read. Connected components in the resulting graph do not change.

The second observation is that in our problem, we never remove (arbitrary) nodes from the overlap graph. This simplifies data management as we can adopt tested data structures such as disjoint-set to identify connected components [14].

In our approach, outlined in the right-most panel of Figure 1, we exploit both properties at the same time: we first provide efficient strategy to identify redundant reads in a stream, and then use a variant of union-find to track connected components. In the process, we provide feedback information to the overlapper to further improve end-to-end performance on the entire workflow.

### 3.1 Finding transitive nodes

The first step in our approach is to decide whether an incoming read is redundant, and hence can be removed from further processing. Let us consider again example reads in Figure 2a, and for the purpose of presentation, let us assume that *r*_*x*_ ≺ *r*_*y*_ ≺ *r*_*i*_, *r*_*j*_, *r*_*k*_. Depending on which of the reads *r*_*i*_, *r*_*j*_, *r*_*k*_ arrives last, we have one of three possible overlap graphs as shown in Figure 2b. Here, read *r*_*i*_ is represented by vertex *v*_*i*_, and shaded nodes correspond to the last arriving read. Any redundant read, in our case read *r*_*j*_, shares suffix-prefix overlap with at least two other reads, in our case *r*_*i*_ and *r*_*k*_. Now let us consider an induced sub-graph over the redundant vertex (i.e., vertex corresponding to the redundant read, in our case *v*_*j*_) and any two of its adjacent vertices that are also adjacent to each other. In this sub-graph, one of the nodes must have two outgoing edges, one must have two incoming edges, and one must have one incoming and one outgoing edge (this node must correspond to a redundant read). Back to Figure 2b, we can see that node *v*_*j*_ is redundant irrespective of the order in which reads are processed due to the transitive edge (*v*_*k*_, *v*_*i*_). We will call nodes that introduce transitive edges between two other nodes *transitive*, and since they correspond to redundant reads, we will eliminate them from processing.

For each incoming read, we now want to identify if it introduces any transitive nodes. This can be done by considering all possible triples it forms with its adjacent nodes. For instance, to find the transitive nodes introduced by the arriving node *v*_*i*_ (i.e., read *r*_*i*_) in the first example in Figure 2b, we need to check if it has a pair of incoming neighbors *u*, *w* ∊ *N*^−^(*v*_*i*_) that share an edge (*u*, *w*) or (*w*, *u*). In the second example, to find the transitive nodes introduced by the arriving node, we check if it has an incoming neighbor *u* ∊ *N*^−^(*v*_*j*_) and outgoing neighbor *w* ∊ *N*^+^(*v*_*j*_) that share an edge (*u*, *w*). Finally, to find the transitive nodes introduced by the arriving node *v*_*k*_, we need to check if it has a pair of outgoing neighbors *u*, *w* ∊ *N*^−^(*v*_*k*_) that share an edge (*u*, *w*) or (*w*, *u*).

In general, we notice that there are three ways an incoming read can introduce transitive nodes. However, because we do not know which case we are dealing with, we need to consider all three. This intuition is captured in Algorithm 1.

For each incoming read *v*, the input to the algorithm are all overlaps with the previously processed nodes (represented by sets *N*^+^(*v*) and *N*^−^(*v*)), and the current irreducible overlap graph *G* (we explain irreducibility below). In lines 3-9, we check if v has a pair of outgoing neighbors *u*, *w* ∊ *N*^+^(*v*) that share an edge (*u*, *w*) ∊ *E* or (*w*, *u*) ∊ *E*. This scenario corresponds to the last case in Figure 2b. In lines 10-16, we check if v has a pair of incoming neighbors *u*, *w* ∊ *N*^−^(*v*) that share an edge (*u*, *w*) ∊ *E* or (*w*, *u*) ∊ *E*. This scenario corresponds to the first case in the Figure 2b. Finally, in lines 17-21, we check if v has a pair of neighbors *u*, *w* where *u* ∊ *N*^+^(*v*) and *w* ∊ *N*^−^(*v*) share an edge (*w*, *u*) ∊ *E*. This scenario corresponds to the middle case in the Figure 2b.

**Algorithm 1.**
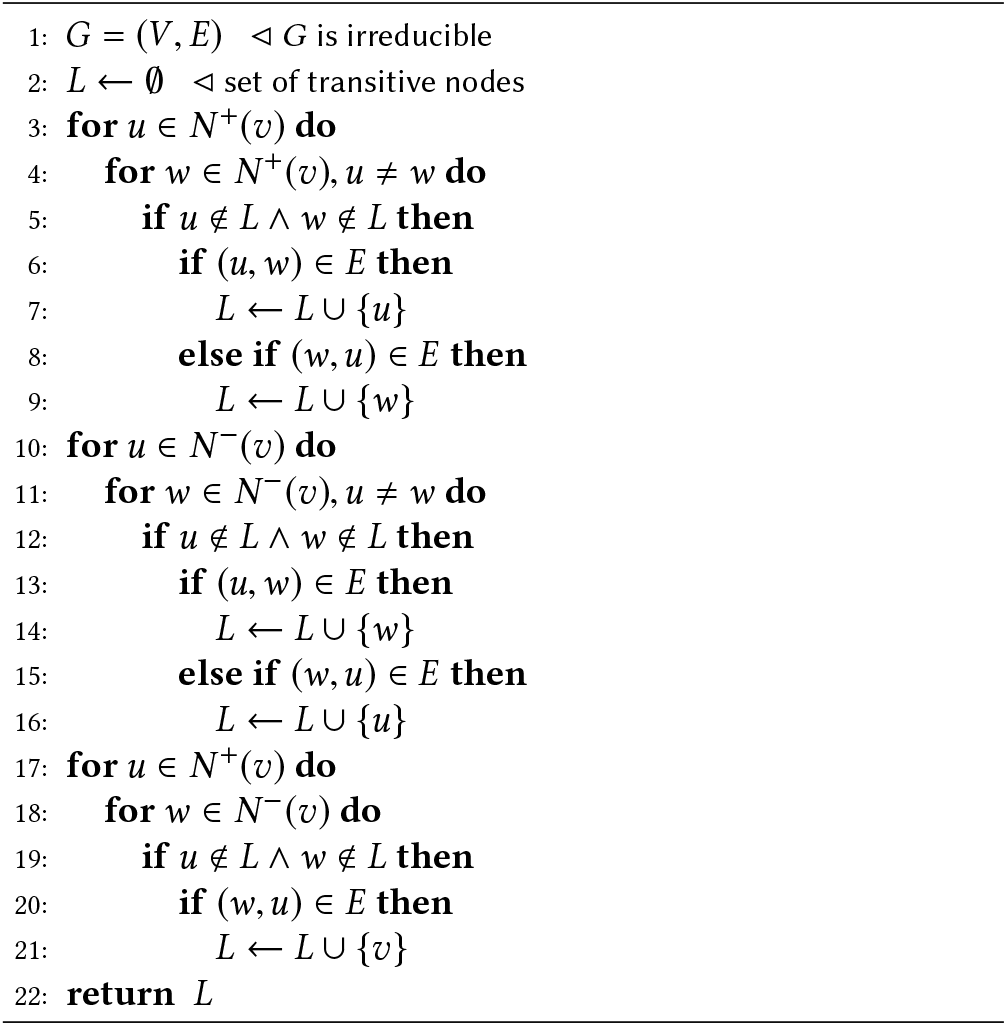
FindTransitive(*G*,*v*, *N*^+^(*v*), *N*^−^(*v*))

Although the processing steps in Algorithm 1 are simple, the entire procedure can be computationally expensive for a mobile system and streaming regime (keeping in mind that it is executed for each incoming read). Because we are searching for edges between incoming neighbors, edges between outgoing neighbors, and edges between incoming and outgoing neighbors, the process requires Θ (|*N*^−^(*v*_*i*_)| + |*N*^+^(*v*_*i*_)|)^2^ edge queries. This is problematic, as *v*_*i*_ may have a high degree, and each edge query can be expensive depending on how *G* is stored in memory. However, we can make certain guarantees about our incoming node degree as long as G^*i*−1^ is *irreducible*. Here, an irreducible graph is an overlap graph that does not contain transitive nodes.

#### Cost of handling transitive nodes

In a streaming regime, we can maintain irreducibility by eliminating transitive nodes the moment they are introduced by an incoming node. Maintaining irreducibility will not require any additional computational effort. However, it will necessitate a dedicated approach to maintain a coherent list of connected components.

Recall that by definition, an overlapper detects an edge (*u*, *v*) if there is a significant overlap between the reads corresponding to *u* and *v*. Here, a significant overlap indicates the reads corresponding to *u* and *v* have been derived from neighboring portions of the underlying genome. If we have an irreducible overlap graph, we are guaranteed that for an incoming read *v*_*i*_ |*N*^−^(*u*_*i*_)| + |*N*^+^(*v*_*i*_)| ≤ 4 with |*N*^−^(*v*_*i*_)| ≤ 2 and |*N*^+^(*v*_*i*_)| ≤ 2. This can be shown through contradiction: Suppose we have an incoming read *r*_*i*_ and an irreducible graph G^*i-1*^. Now, suppose |*N*^+^(*v*_*i*_)| = 3 (or |*N*^−^(*v*_*i*_)| = 3. both work). By definition, the three reads in *N*^+^(*v*_*i*_) that overlap with *r*_*i*_ belong to the same portion of the genome as **r*_*i*_*. This is because their prefixes overlap with *r*_*i*_. Since all three of the reads belong to the same portion of the genome, then they must also overlap with one another. Since they all overlap with one another, at least one of the reads is either contained within another read or contained within the overlap of the other two reads. This means that at least one of the reads is transitive which contradicts that *G*^*i*−1^ is irreducible. This contradiction shows that both |*N*^+^(*v*_*i*_)| and |*N*^−^(*v*_*i*_)| ≤ 2 and therefore |*N*^−^(*v*_*i*_)| + |*N*^+^(*v*_*i*_)| ≤ 4 so long as *G*^*i*−1^ is irreducible. Notice that this is demonstrated in Figure 2b.

We recognize that the above reasoning holds for the case where overlap between reads indicates that they are derived from the neighboring portion of a genome. This assumption is not always true in the real world (e.g., due to repeats in a genome). However, as we show later in Section 3.4, our method still performs well even when the assumption of irreducibility is violated.

As we already discussed, we can guarantee *G*^*i*−1^ is irreducible by removing transitive nodes with each incoming read. We will call removing transitive nodes *transitive closures*.

### 3.2 Maintaining graph and components

Given an efficient routine to identify transitive nodes in the stream, we need a way to maintain both graph structure and the corresponding connected components. Our solution must also be able to handle deletions of transitive nodes.

To maintain graph *G*^*i*^, we use a simple adjacency list built on top of a hash table. For each vertex we maintain a list of its incoming and outgoing neighbors, and the neighbor lists are kept in a hash table with key derived from vertex identifier. In this way, we compensate for the fact that the size of the input stream is not known in advance. This solution is practical and efficient, and takes into account small expected size of the neighbor lists.

To store and track connected components, we use a variant of union-find data structure (UF). Specifically, we represent UF as a set of key-value pairs 〈*v*_*r*_, *v*_*c*_〉, where *v*_*r*_ is some node mapped to a component represented by *v*_*c*_ (i.e., *v*_*c*_ is the root of the component). Here, set is again implemented over a hash table, to account for the fact that the size of UF will be changing dynamically.

**Algorithm 2.**
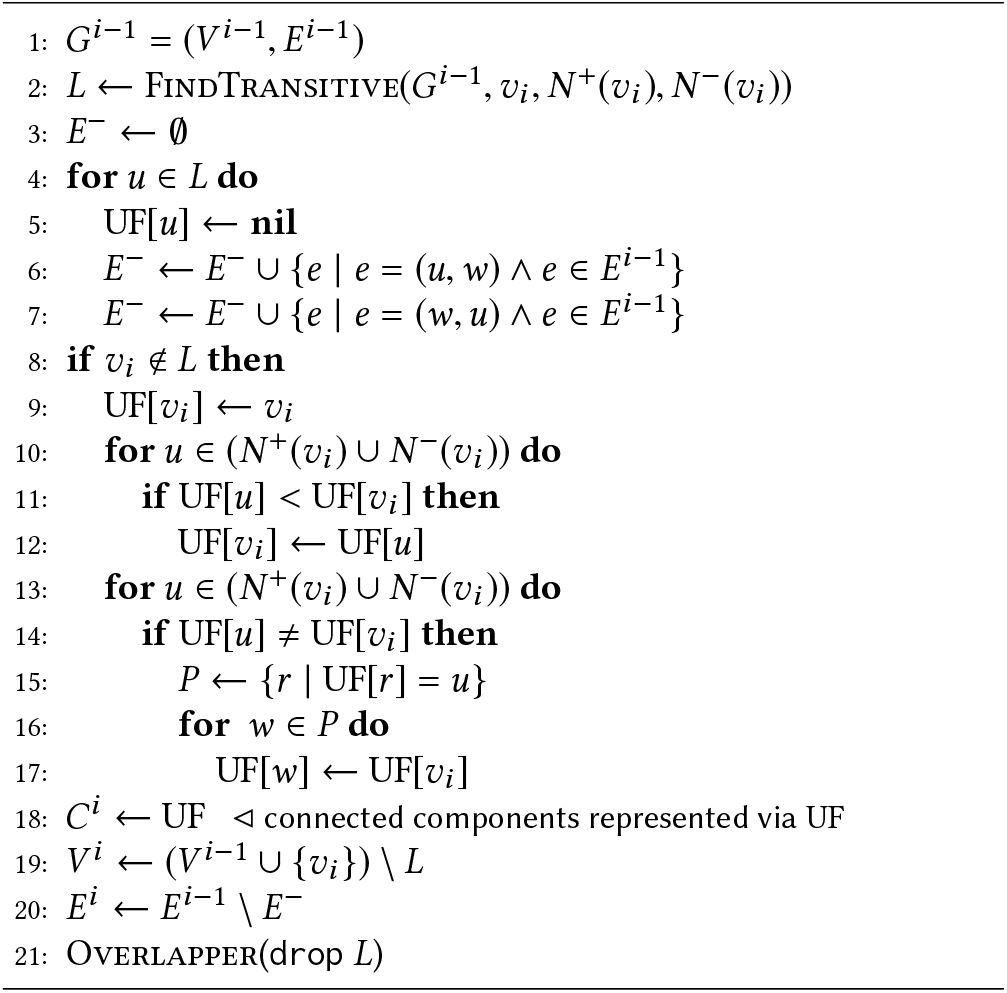
StreamCC(*G*^*i*−1^, UF,*v*_*i*_ *N*^+^(*v*_*i*_), *N*^−^(*v*_*i*_))

### 3.3 Maintaining connected components over stream

Having all ingredients in place, we summarize our method for maintaining connected components in Algorithm 2. In the first step, we identify all transitive nodes that are safe to remove using our FindTransitive routine (line 2). For each transitive node, we first remove its corresponding entry from UF structure (line 5), and we identify all associated edges to remove from *G* (lines 6 and 7). Because we are using hash tables for both UF and *E*, both of these operations can be done in amortized *O*(|*V*|). Since *G* is reduced by transitive closures, we expect |*V*| to be a small fraction of all processed nodes (which we confirmed via experimental results). If the incoming node vi is not in the list of nodes to remove, we proceed with insertion in lines 8-17 (explained below). Finally, in lines 18-21, we remove all edges associated with transitive nodes along with the transitive nodes and reads themselves.

One significant advantage of our approach is that it can provide feedback to the overlapper (line 21). Since transitive nodes correspond to redundant reads, they may be offloaded (or dropped) to a persistent storage, and processed later as needed, thus saving memory. Moreover, by maintaining only non-redundant reads, an overlapper may improve its performance as well (although this could potentially come at the price of missing some of the overlaps).

#### Inserting nodes

To insert a node *v*_*i*_ (lines 8-17), we process how *v*_*i*_ affects UF (lines 9-17). Node *v*_*i*_ can form its own component (if it has no neighbors), join a component (if its neighbors all belong to the same component), or merge components (ifits neighbors belong to different components). First, we assume that *v*_*i*_ is forming its own component in line 11. Then, we handle the situation where it joins a component or merges components. If *v*_*i*_ has neighbors, then we identify the root of the component to which *v*_*i*_ belongs to (lines 10-12). We note that because we are incrementally maintaining UF with each incoming read, the resulting structure always has a depth of one (all nodes are connected to the root). Therefore, the cost of this step is *O* (|*N*^−^(*v*_*i*_) ∪ *N*^+^(*v*_*i*_)|), which is constant when *G* is irreducible.

In lines 13-17, we handle the situation where components are merged. We do this by checking if any of *v*_*i*_’s neighboring nodes have a different root (line 14). When this is the case, we reassign all of the nodes 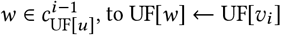. As a result, neighbors of *v*_*i*_ end up having the same component mapping as *v*_*i*_. Note that gathering component members in line 15 requires searching through UF, and hence takes *O*(|*V*|) time. However, this cost gets amortized over the course of execution as shown in experimental results.

### 3.4 Tuning to real world data

So far we have been working under the assumption that an overlapper correctly identifies suffix-prefix overlaps between reads coming from the neighboring parts of a genome. Although this assumption is helpful in our understanding of transitive nodes, it is not entirely realistic. This is because DNA reads, especially in MinION sequencing, are error prone and typical genomes are repetitive. Consequently, an overlapper may either miss overlap between reads (e.g., due to sequencing error) or may detect spurious overlaps (e.g., due to repetition where similar reads come from different regions of a genome).

If an overlapper detects an overlap between reads that do not belong to neighboring portions of the genome, then this is a false positive. The connected components of the overlap graph that contains many false positives may not be a robust representations of OTUs.

This is because false positive edges may link reads from distinct OTUs, and merge their respective components. If an overlapper misses an overlap between two reads that belong to neighboring portions of a genome, then this is a false negative. This can cause two components that are supposed to be connected to be disjoint. Disconnected components may also misrepresent the true OTUs, leading to incorrect assessment of the number of OTUs.

In Figure 3, we show a hypothetical scenario that demonstrates the impact of false positive/negative edges. In this scenario, node *u* is transitive, and therefore it will be removed by transitive closure. However, in the presence of false positive/negative edges, removing a transitive node may disconnect our graph. To see why, observe that nodes *w* and *v*_*i*_ are connected to *u*, and *x* and *y* are connected to *u*, and yet these two pairs are not connected to each other. This could have happened because overlapper missed edges (*u*, *x*) and (*v*_*i*_, *y*) or the overlapper incorrectly identified edges (*x*, *u*) and (*u*, *y*). In either case, removing *u* will disconnect the graph. We will call nodes that disconnect components upon removal *articulation points*.

**Figure 3:**
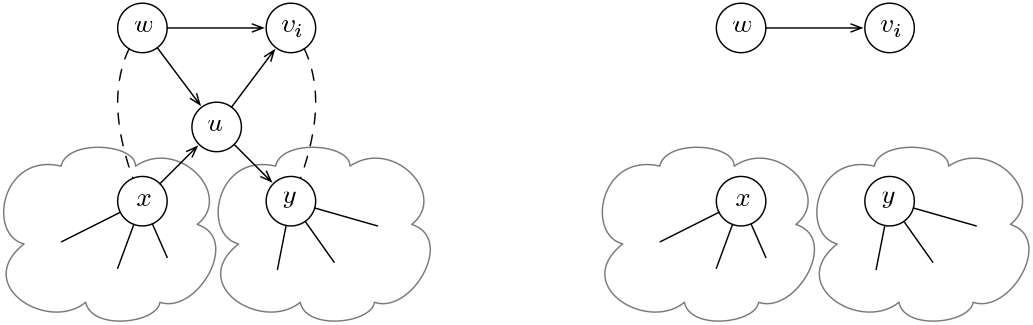
Impact of false positive and false negative edges on connected components discovery when using transitive closure. Left: vertex *m* is added to the graph with single component consisting of vertices *{w, u, x, y}. We* consider two cases where edges *(w, x)* and *y)* are incorrectly missing, or edges *(x, u)* and *(u, y)* are incorrectly introduced. Right: After performing transitive closure on *u,* in both cases the graph will consist of three connected components instead of one component.

#### Protecting articulation points

When transitive nodes are articulation points, we know this is due to an overlapper error. However, we can not conclude whether this is due to false positive or false negative edges. Because of this uncertainty, we cannot assume that articulation points correctly provide redundant information, and therefore should not remove them. Moreover, disconnecting components would require recomputing all components from scratch thus degrading computational performance. Furthermore, to even determine whether a component has become disconnected is a challenging problem in its own right [17]. Hence, in our approach we decided to protect articulation points from being removed.

Let nodes corresponding to the reads that contain the redundant read be *anchor nodes*. For example, in Figure 3, *u* is transitive while *v*_*i*_ and *w* are anchor nodes. Instead of checking whether a node is an articulation point, we will impose a more strict and inexpensive to compute criteria: A transitive node must have *anchored* neighbors for it to be removed. A node is anchored if it shares an edge with an anchor node. If all of a transitive node’s neighbors are anchored, then we can safely remove the transitive node. This is because a transitive node cannot be an articulation point if its neighbors are anchored. Anchor nodes must belong to the same component as the transitive node. If all of the transitive node’s neighbors are anchored, then they all have at least two points of connection to the rest of the component: one point is with the transitive node and another point is with at least one of the anchored nodes. This allows us to safely remove the transitive node because the neighbors still maintain at least one point of connection to the rest of the component.

We can modify Algorithm 2 to enforce that we do not remove transitive nodes that may be articulation points. To do this, it is sufficient that we remove from set *L* any node whose neighbors are not all anchored. For each node *u* ∊ *L*, this necessary condition can be checked in *O* (|*N*^−^(*u*)| + |*N*^+^(*u*)|) time because each neighbor of *u* requires a constant number of edge queries to see if it is anchored. Hence, this modification imposes only slight overhead compared to the original algorithm.

#### Effects of preserving transitive nodes

We will refer to an overlap graph that has transitive nodes that are not articulation points removed as a *reduced* graph. In a reduced graph, we are guaranteed to not disconnect an existing component. However, in the following example, we illustrate how removing transitive nodes may still result in a fragmented component in the future. Suppose that nodes *u*, *v*_*i*_ and *w* precede nodes *x* and *y* in Figure 3 (*u*, *m*, *w* ≺ *x*, *y*). Since *x* and *y* have not been added to the graph yet, node *u* is not an articulation point and hence is removed by transitive closure. When *x* and *y* are added to the graph, they must end up in different components from *w* and *v*_*i*_ since *u* is missing. Although this may affect the component count, in the experimental results, we show that connected components are still robust representations of OTUs.

Another observation is that because we cannot remove all transitive nodes, we will not have the same degree guarantees as in an irreducible overlap graph. For example, in Figure 3, we cannot remove *u* due to it being an articulation point. Later in the process, we may receive a read *r*_*j*_, that has specifically sufRx-prefix overlaps with the reads corresponding to *w* and *x* but also *u*. In such case, the out degree of *v*_*j*_ will be three, which violates our previously established degree constraints in irreducible graphs.

Fortunately, in practice, we observe that incoming nodes in the stream follow an exponential degree distribution, which is a side effect of transitive closures (see experimental results). If we assume an exponential distribution, then we can still guarantee that the expected degree of a node will be constant. Suppose an incoming node *v*_*i*_ has degree of *k* with probability *p*_*k*_ where *p*_*k*_ = (1 − *e*^−1/*k*^)e^−*K*/*k*^ and *k* is some constant [23]. Then the expected average degree of an infinite stream is:

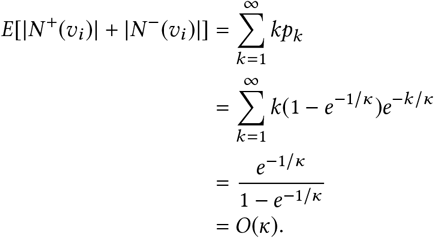

Since the average degree is bounded by *κ*, as long as *κ* is a small constant, maintaining transitive closures on a reduced graph will still perform similarly to an irreducible overlap graph. In our experimental results, we found that *κ* never exceeds three.

## 4 Experimental Results

To validate our proposed approach we implemented Algorithms 1 and 2, including the extensions from Section 3.4, in a standalone C++ application (the code is open source and available from [33]). We performed a set of experiments using synthetic as well as the actual, publicly available, GridlON data. In our experiments, we focused on performance (e.g., run time and memory use) as well as correctness characteristics (e.g., connected components and OTUs recovery and convergence) taking into account properties of the overlapper. We also note that all experiments that involved random sampling or shuffling of the data were repeated multiple times, and we did not observe significant difference from the results reported below.

### 4.1 Test data

To prepare benchmark data, we started from the `ERR3152364` dataset publicly available from the European Nucleotide Archive [3]. This dataset has been generated using the ONT GridlON sequencer, which is a scaled-up version of the portable MinION platform. We selected this particular dataset since it represents metagenomic sequencing of a mock community with known microbial composition. Specifically, the dataset is based on the Zymo Community Standards that comprises five Gram-positive bacteria, three Gram-negative bacteria, all eight organisms in the same abundance, and two types of yeast that make 4% of the community (more details regarding this particular dataset and how sequencing has been performed can be found in [24]).

To annotate the reads, i.e., assign them to one of the component organisms in the mock community, we performed read mapping using the latest blast tool and reference genomes provided by Zymo [2, 35] (maker of the mock community). For each mapped read, we selected its target OTU based on the best mapping score (i.e., to which reference genome it mapped the best). We disregarded reads with low mapping scores, and reads that did not map uniquely. Furthermore, we eliminated veiy few reads mapping to yeast genome since they made less than 4% of all reads. The resulting dataset consists of 2,998,869 classified (i.e., assigned to an OTU) reads. We will refer to the resulting dataset as `ERR3152364`.

We used annotated `ERR3152364` dataset to simulate artificial but realistic reads using the `NanoSim tool` [32]. Starting from the actual GridlON reads and the corresponding reference genomes, `NanoSim` first builds a statistical model of the GridlON sequencer, and then uses this model to derive new reads from the reference genomes.

Since `NanoSim` reports from which position in the genome each read has been derived, we can use this information to create a perfect overlapper, i.e., an overlapper in which no false positive or false negative edges are created. This enables us also to control for overlapper performance (e.g., precision and sensitivity) when assessing performance of our algorithms. We refer to the resulting dataset as `Sim`. The dataset consists of 200,000 reads, where each of the eight reference genomes is represented by 25,000 reads.

For reproducibility purposes, we provide all details regarding data preparation in the accompanying web page [33].

### 4.2 Overlapper

Detection of prefix-suffix overlaps in DNA reads is extensively studied topic, especially in the context of long reads such as those produced by MinION platform. However, although there are several overlappers available (e.g., `daligner` [21], `minimap2` [15], `MHAP` [4], `ELaSTIC` [34]), these tools do not provide any formal guarantees with respect to the quality and correctness of the discovered overlaps. Moreover, they currently are not meant to work in a streaming regime, and are not designed to incorporate potential feedback provided by our algorithms (e.g., line 21 in Algorithm 1). We note that right now we are in the process of developing a custom streaming-based overlap tool specifically for mobile DNA analytics workflows, which will address the above shortcomings, and will complement the solutions presented in this paper. Taking all of that into account, in order to test our solution while controlling for overlapper quality and performance, we decided to simulate overlap graph streaming based on overlaps detected via batch processing.

To simulate the process for the `ERR3152364` dataset, we leveraged directly information provided by the blast tool when performing reads assignment to OTUs (as explained in the previous section). Specifically, we assumed that two reads overlap if they are mapped by blast to the same portions of the underlying reference genome such that they have a suffix-prefix overlap of at least size 1,000 nucleotides/characters (based on [20] we consider such overlap length significant). To simulate an overlapper for the `Sim` dataset, we used a similar approach. However, instead of using `blast`, we directly used the exact mapping information provided by `NanoSim` (as explained earlier). In both cases, we obtained a complete overlap graph *a priori,* and hence we were able to directly emulate a stream of reads *R* and their neighborhoods *N*^+^ and *N*^−^.

### 4.3 Connected components recovery

In the first experiment, we tested how maintaining transitive closures, hence eliminating redundant reads, affects our ability to recover connected components and their corresponding OTUs. To do this, we constructed a series of overlap graphs over the Sim dataset, in each case randomly removing a fraction of edges to introduce false negatives. For each such created overlap graph, we simulated a streaming process and used our software to identify connected components from the stream. To obtain the actual number of connected components in a given overlap graph, we used standard union-find algorithm on the graph without streaming. We note that in the ideal scenario, both methods should be identifying eight connected components, each corresponding to one OTU in the dataset. The results of this experiment are summarized in Figure 4. Here we use Actual to denote the number of connected components that are identifiable in the graph, and Transitive to denote the number of connected components recovered when the graph is processed using our method.

**Figure 4:**
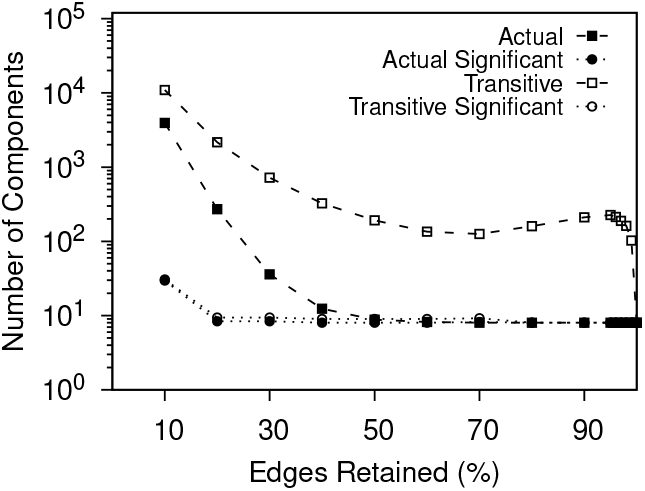
Number of identified connected components depending on the percentage of edges retained in the `Sim` overlap graph that has eight components (*y*-axis is in log scale). Actual means components identified using union-find on the entire graph, Transitive means components identified using our method. Significant means components with more than three nodes.

From the figure we can make several observations. First, in order to correctly recover all eight OTUs the overlap graph must contain at least 50% of edges. When edge retention is below 50%, the missing edges cause the graph to become highly disconnected and hence components no longer uniquely identify OTUs. Second, irrespective of the edge retention, transitive closures introduce exponentially more connected components than there are in the unprocessed graph. This is not entirely surprising considering our previous discussion on how missing edges can lead to disconnected nodes with transitive closures. As edge retention increases, so does expected node degree of the nodes in the underlying graph. This in turn introduces more redundancies therefore increasing the number of transitive closures performed. In our tests, the impact of transitive closures become less pronounced only after exceeding 95% edge retention resulting in sharp dip visible in Figure 4.

While these results may suggest poor performance of our method, the picture drastically changes if we consider only connected components with more than one node. In Figure 5, we inspect the component size distribution of components recovered in the streamed and actual graph. As edge retention increases, we observe that there are eight distinct components along with primarily singleton components. This clear gap in component sizes allows us to easily distinguish significant components that correspond to the target OTUs. If we assume that significant components are components of at least size three (denoted by Significant in Figure 4), then from Figure 4, we can see that we quickly converge to the desired number of components representing OTUs in both our streamed and actual graph. We note that we observe the same behavior in real world dataset `ERR3152364`.

**Figure 5:**
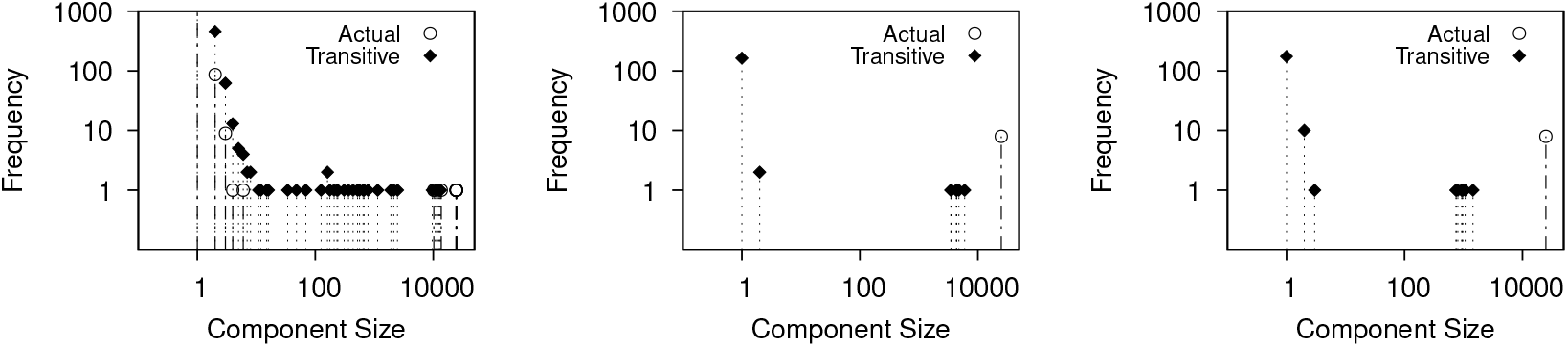
Frequency of components of given size when 10% (left plot), 50% (middle plot) and 90% (right plot) of the edges in the `Sim` overlap graph are retained (both axes are in log scale). As edge retention increases, there are eight large components along with primarily singleton components.

### 4.4 Performance and memory characteristics

In the second set of experiments, we accessed performance characteristics of our algorithm in terms of runtime and memory use. As discussed earlier, the cost of performing transitive closures and incrementally maintaining connected components depends on the average degree of each incoming node, irrespective of the performance of the overlapper. In Figure 6, we plot the incoming node degree distribution against the fitted exponential distribution when streaming the Sim dataset. From the figure, we can see that the exponential distribution indeed closely fits our data for various levels of edge retention. Moreover, K (parameter of the underlying exponential distribution) consistently remains a small constant. This confirms that transitive closures can be performed in amortized constant time. We note that we observe the same behavior in real world dataset `ERR3152364`.

**Figure 6:**
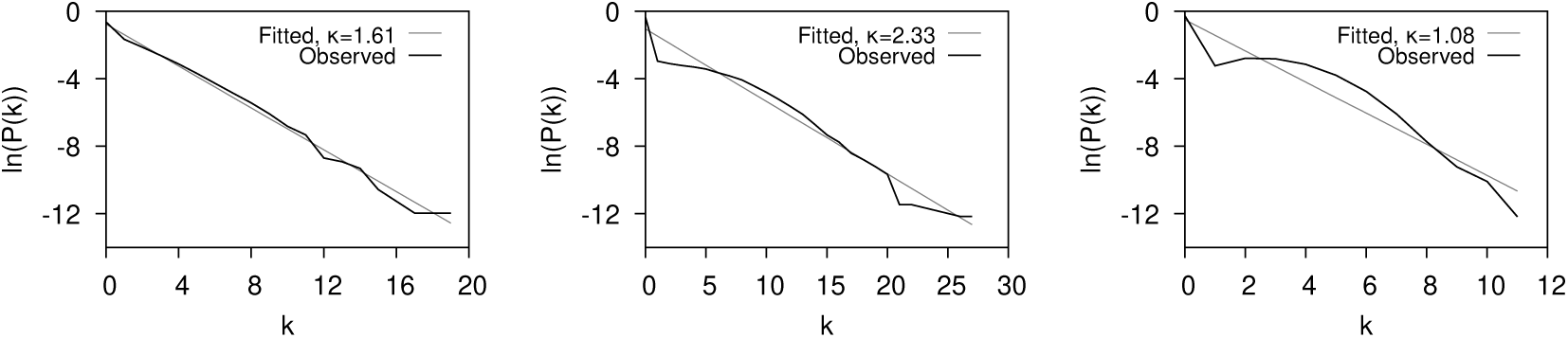
Node degree distribution observed when streaming `Sim` dataset and corresponding fitted exponential distribution, when 10% (left plot, *k = 1.61*, *R*^2^ = 0.99), 50% (middle plot, *k* = 2.33, *R*^2^ = 0.96) and 90% (right plot, *k = 1.08*, *R*^2^ = 0.92) of the edges are retained.

In Figure 7, we show how eliminating redundant reads through transitive closures affects storage rates throughout the streaming process. Here, storage rate refers to the fraction of reads as well as their corresponding graph nodes that we preserve in the main memory. From this figure, we can see that as edge retention improves (i.e., we have fewer false negatives), we store a smaller percentage of nodes. This makes sense intuitively because increased edge retention introduces more redundancies, which lead to more transitive closures. The same reasoning can be applied to explain why storage use improves over time. Specifically, as more nodes are introduced, more overlaps are being detected and hence more redundancies are discovered and eliminated via transitive closures.

**Figure 7:**
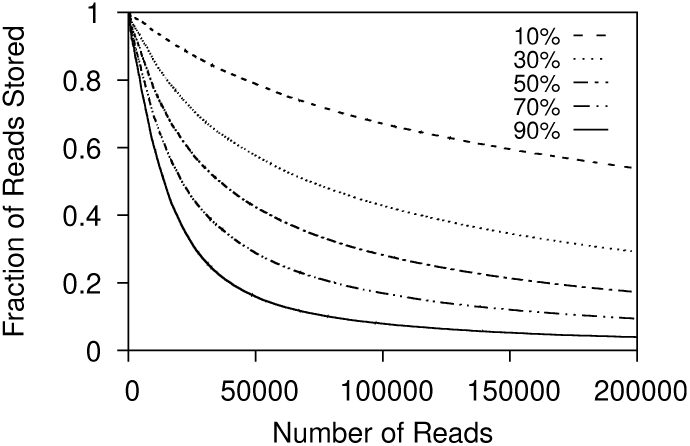
Fraction of nodes maintained in the memory as a function of the number of reads processed for cases where different percentage of edges are retained in the `Sim` dataset.

To assess the cost of incrementally maintaining connected components, as discussed in Section 3.3, we show the effect of incoming reads on component formation in Figure 8. Initially, when nodes enter the graph, they form singletons. Then, the components grow larger and begin to merge thus reducing the number of components. Finally, the number levels off and remains constant. This indicates that as components get larger, merging becomes less frequent, supporting our claim that the costly task of merging components gets amortized over the course of execution.

**Figure 8:**
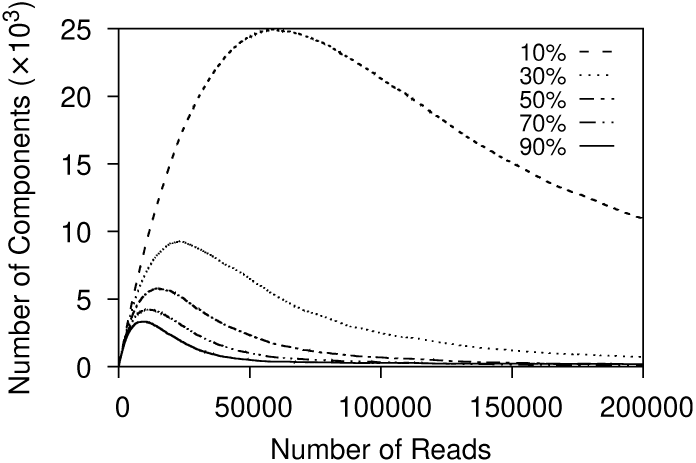
Number of connected components identified as a function of the number of reads processed for cases where different percentage of edges are retained in the `Sim` dataset.

### 4.5 Real world data

In the final set of experiments, we assessed how effective our proposed solution is in recovering OTUs in real world data. We classified all reads in the `ERR3152364` dataset as discussed earlier, and used the resulting classification to assign them to OTUs. To obtain the actual number of connected components in the `ERR3152364` overlap graph, we used again standard union-find algorithm. The algorithm returned ten connected components and nine of them were significant instead of the expected eight. This discrepancy between the number of connected components and the number of OTUs is explained by the complexity of one of the reference genomes. Specifically, the genome of *Salmonella* is highly repetitive, which causes repetitive reads to cluster in one region of the genome thus not providing sufficient information to connect other regions.

We simulated the streaming process of the `ERR3152364` overlap graph, and recorded the same statistics as for the simulated data. Our method was able to recover all nine significant components (and thus their corresponding OTUs) after processing 1,504,145 reads (about half of all reads). In our experiments, we found that the exponential degree distribution was a good fit with *R*^2^ = 0.81 and κ = 2.4. This confirms that the assumption of amortized constant time for performing transitive closures holds for the real world dataset. Finally, throughout processing, the maximum number of nodes stored was 28,524 (less than 1% of total reads processed) clearly demonstrating effectiveness of maintaining transitive closures.

## 5 Related Work

The idea of using transitive closure to reduce computational complexity has been used previously in the context of DNA assembly, where DNA reads are pieced together to reconstruct longer DNA strings [26]. For example, in [19], Myers discusses DNA string graphs in which vertices represent prefixes and suffixes of reads, and edges represent non-matching substrings between two overlapping reads. He then shows that transitive edges can be removed without affecting the ability to reconstruct the assembly. While in our approach we essentially exploit the same principle, i.e., redundancy of overlapping reads, our focus is on streaming and not memory prohibitive batch processing.

Disjoint-set forests, also known as union-find, is the most efficient data structure to find connected components in a graph. Because of its popularity and versatility, there are several adaptations of union-find for various computational setups. For example, Isenburg and Shewcuk [11] adapted the union-find algorithm for a streaming 3D grid network to use in image processing, Agarwal et al. considered I/O efficient solutions for terrain analysis [1], and Simsiri et al. studied work-efficient parallel adaptations of union-find for incremental graph connectivity [29]. Laura and Santaroni introduced the first semi-streaming algorithm that makes a few passes to find strongly connected components in a directed graph [14]. These methods, however, are primarily focused on general streams where graph nodes and edges can be inserted or removed at any point of time. Moreover, they assume that the executing environment has significant main memory available. In our case, the problem has a slightly different flavor. On the one hand, the stream is easier to handle because we consider only node insertions and specific node removals. Due to the nature of metagenomic reads, we can also expect bounded number of edges introduced with every node insertion. On the other hand, in any mobile setup we have very limited access to the main memory and computational power (typically, available memory is around 4GB). Consequently, our primary focus is on maintaining minimal memory footprint, while delivering the desired statistics.

## 6 Conclusion

The growing popularity and rapid adoption of portable DNA sequencing platforms necessitates the development of new computational strategies to enable *in situ* DNA analytics. In this work, we introduce OTUs identification method tailored for low-memory mobile devices. The method can be used to accelerate end-to-end mobile execution of multiple types of DNA analysis, including assembly, metagenomic classification, etc.

The key element of our solution is memory efficient handling of connected components emerging in streams of DNA reads and their overlap graphs. Through formal and experimental analysis, we show that if the degree of nodes in the streamed overlap graph follows exponential distribution (which is the case in real-world instances) our method has minimal computational cost of handling incoming reads. Consequently, the method is ideal for mobile computing.

While our proposed solution addresses the question of how to identify components (or OTUs) in a stream, it is based on the assumption that the processing pipeline includes an overlapper that is able to work efficiently over the streamed DNA reads. While such overlappers are not yet readily available, we are currently investigating an adaptive overlapper operating directly on the raw signals produced by MinION sequencer (i.e., bypassing basecalling stage). Both components are part of SMARTEn [28], our broader effort in mobile DNA processing.

## 7 Acknowledgement

We would like to acknowledge the National Science Foundation for supporting this work under the grant CNS-1910193.

